# Muscone-specific olfactory protein reveals the putative scent-marking pheromone in the Sunda pangolin (*Manis javanica*)

**DOI:** 10.1101/2024.06.04.597258

**Authors:** Zhongbo Yu, Tao Meng, Luyao Yu, Yichen Zhou, Tengcheng Que, Meihong He, Haijing Wang, Yingjiao Li, Liling Liu, Wenjian Liu, Yinliang Wang, Bingzhong Ren

## Abstract

The Sunda pangolin (*Manis javanica*) is a burrowing and nocturnal animal, and they have poor vision, thus, intraspecies communication relies on olfaction, such as mating, warning, and scent-marking. However, the intraspecies pheromone in pangolins remains unknown. In this study, all the odorant-binding proteins in Sunda pangolins were functionally expressed, and they were screened against a panel of 32 volatiles that were derived from the pangolin’s urine, feces, and anal gland secretions. Reverse chemical ecology identified that *M. javanica* odorant-binding protein 3 (MjavOBP3) possesses the highest binding affinity to muscone. A subsequent behavior-tracking assay showed that only males can sense muscone; thus, we hypothesize that muscone is a male-specific scent-marking pheromone. Meanwhile, the structural study showed that Tyr117 contributes the most to muscone’s binding, which was further validated by site-directed mutagenesis. The findings clarify the scent-marking mechanism in pangolins, and muscone could potentially be used to support the monitoring and conservation of this endangered animal.

**Author Summary:** The Sunda pangolin is an endangered mammal that is native to Southeast Asia and is threatened due to its economic value. They are cave-living and nocturnal, poor vision; thus, their intraspecies communication is highly reliant on olfaction. Although they are generally solitary, they have been observed to have some social aspects in the wild, such as breeding and territorial behaviors, which are mediated by scents. However, no previous study has investigated the type of pheromones and how they are detected. Using the reverse chemical ecology approach, MjavOBP3 was found to bind to muscone with high affinity, and behavior-tracking assay was performed under well-controlled artificial rearing conditions, which showed only male pangolins can recognize muscone, suggesting its potential male-specific pheromone role.

## Introduction

Scent-marking is one of the most conspicuous behaviors of mammals and other terrestrial vertebrates. It has long intrigued researchers from various disciplines [1]. Although the chemical composition, source, and behavioral effect of scent-marking are highly diverse among different taxa, it has been suggested to have four main functions: 1) attracting females to increase mating and reproductive success [2]; 2) advertising males’ territory ownership and social dominance status [3]; 3) marking of food resources to increase foraging efficiency [4]; and 4) individual recognition among different species and populations [5]. In some cases, scent marking not only advertises the existence of the scent owners but also includes information about the individual’s health, infections, age, sex, and mating state [5–7]. The source of the scent marking that is used by mammals is usually urine, feces, and glandular secretions [8]. For instance, Siberian wolves (*Canis lupus*) deposit feces at crossroads to increase the effectiveness of territory maintenance [9, 10], and female Asian elephants (*Elephas maximus*) use urine to signal males of their readiness to mate [11]. Anal gland secretions are also used by some Carnivora, such as ferrets (*Mustela putorius furo*), meerkats (*Suricata suricatta*), domestic cats (*Felis catus*), and giant panda (*Ailuropoda melanoleuca*), and the scent that is emitted from their anal gland provides important information for intraspecies individual recognition [12–15].

The Sunda pangolin belongs to Manidae, Pholidota, and is closely related to the family Carnivora. There are only eight extant species in the world [16]. Due to the economic value of their meat, scales (attributed medicinal properties), and skins, enormous numbers of pangolins have been seized and traded in the black market, especially in Southeast Asia [17]. Currently, *M. javanica* is classified as a critically endangered species and is listed in Appendix II of the Convention on International Trade in Endangered Species of Wild Fauna and Flora (CITES) [18]. Malayan pangolins is nocturnal and has a highly specialized diet, foraging only on ants and termites, with genomic evidence showing that their olfactory receptor gene families are significantly expanded, supporting that they have a well-developed olfactory system [19]. Pangolins are generally solitary but they also have home ranges [20], which are more distinguished in males. Male pangolins have been observed to display aggressive behavior toward each other to defend their home ranges, suggesting that they are territorial [21]. Most of the social interactions of *M. javanica* are believed to be scent-based and generally comprised of urine and anal glands secretions [20]; however, well-controlled behavioral experiments on *M. javanica* are still lacking, and our knowledge of the behaviors of *M. javanica* still relies on the observations from zoos and nature reserves.

Consequently, the reverse chemical ecology approach may shed light on the communication behaviors of Sunda pangolin, and it is a simple and effective tool for screening biologically active scents and clarifying their molecular mechanisms [22]. Among the olfactory proteins, odorant binding proteins (OBPs), which belong to the lipocalin family, might be specifically dedicated to pheromonal communication [23]. When compared with insect OBPs, the function of vertebrate OBPs remains less well understood; however, evidence suggests that these proteins might also be involved in scent markings. For instance, three OBPs were identified in the anal sac gland of dogs (*Canis lupus familiaris*) and are involved in the signal release and nasal mucosa reception of scent secretions [24]. Similarly, rodent urine also contains a class of lipocalins that play a key role in individual recognition [25]. Although the genome of *M. javanica* was sequenced many years ago, at present, the functional expression of proteins of this kind has not been studied in this species. Moreover, the behavioral and molecular mechanisms of scent marking in Sunda pangolin remain elusive.

Consequently, in this study, we analyzed the *M. javanica* transcriptome from different tissues, and identified, cloned, and analyzed the expression of all three *M. javanica* OBPs (MjavOBPs). Based on the reverse chemical ecology approach, we screened these OBPs against a panel of volatiles from their urine, feces, and anal gland secretions with the aim to explore the scent-marking-related chemical signals. Additionally, the behavioral effect of the active ligands was evaluated using a behavioral tracking assay. Lastly, the key binding sites of the OBP-ligand complex were identified and verified by molecular modeling to clarify the structural-based selectivity of the MjavOBPs.

## Results

### Candidate volatiles were associated with scent marking

Forty-nine volatiles were identified by a gas chromatography-mass spectrometry (GC-MS) analysis of urine, feces, and anal gland secretions (Fig 1; S1 Table), namely: 21 fatty acids, seven ketones, seven alcohols, four esters, three alkanes, three aromatic compounds, and four other compounds. The top three abundant volatiles in the feces of the males and females were 4-cresol (♂: 40.01 ± 2.7%, ♀: 44.67 ± 4.33%), isovaleric acid (♂: 10.58 ± 0.76%, ♀: 9.01 ± 1.72%), and butanoic acid (♂: 7.05 ± 0.75%, ♀: 10.44 ± 1.24%; Fig 1A and B). Then, 3-methylindole, a common volatile in feces, was detected (♂: 0.66 ± 0.51%, ♀: 4.84 ± 1.55%), and, the ant volatiles dimethyl trisulfide (♂: 2.67 ± 1.64%, ♀: 4.24 ± 1.02%) and farnesene (♂: 0.24 ± 0.01%, ♀: 0.09 ± 0.04%) were also detected, which might be correlated with the pangolin’s foraging habitats. The top three volatiles in the urine of the males and females had minor differences, with hexadecanoic acid (33.63 ± 1.20%), zibeton (22.37 ± 4.97%), and benzaldehyde (15.00 ± 4.03%) being the top three volatiles in males, and those in females were hexadecanoic acid (31.73 ± 7.19%), tetradecanoic acid (22.04 ± 3.05%), and zibeton (12.50 ± 2.29%; Fig 1C and D). Additionally, phenol (14.82 ± 4.06%), 2-methylheptanoic acid (10.76 ± 0.21%), and 2-methyldecanoic acid (10.22 ± 0.62%) were the most abundant volatiles in the anal gland secretions (Fig 1E). Interestingly, muscone, which is the pheromone in musk deer, and 8-cyclohexadecen-1-one, a volatile that has a similar chemical structure to muscone, were detected in the anal secretions.

**Figure 1.**
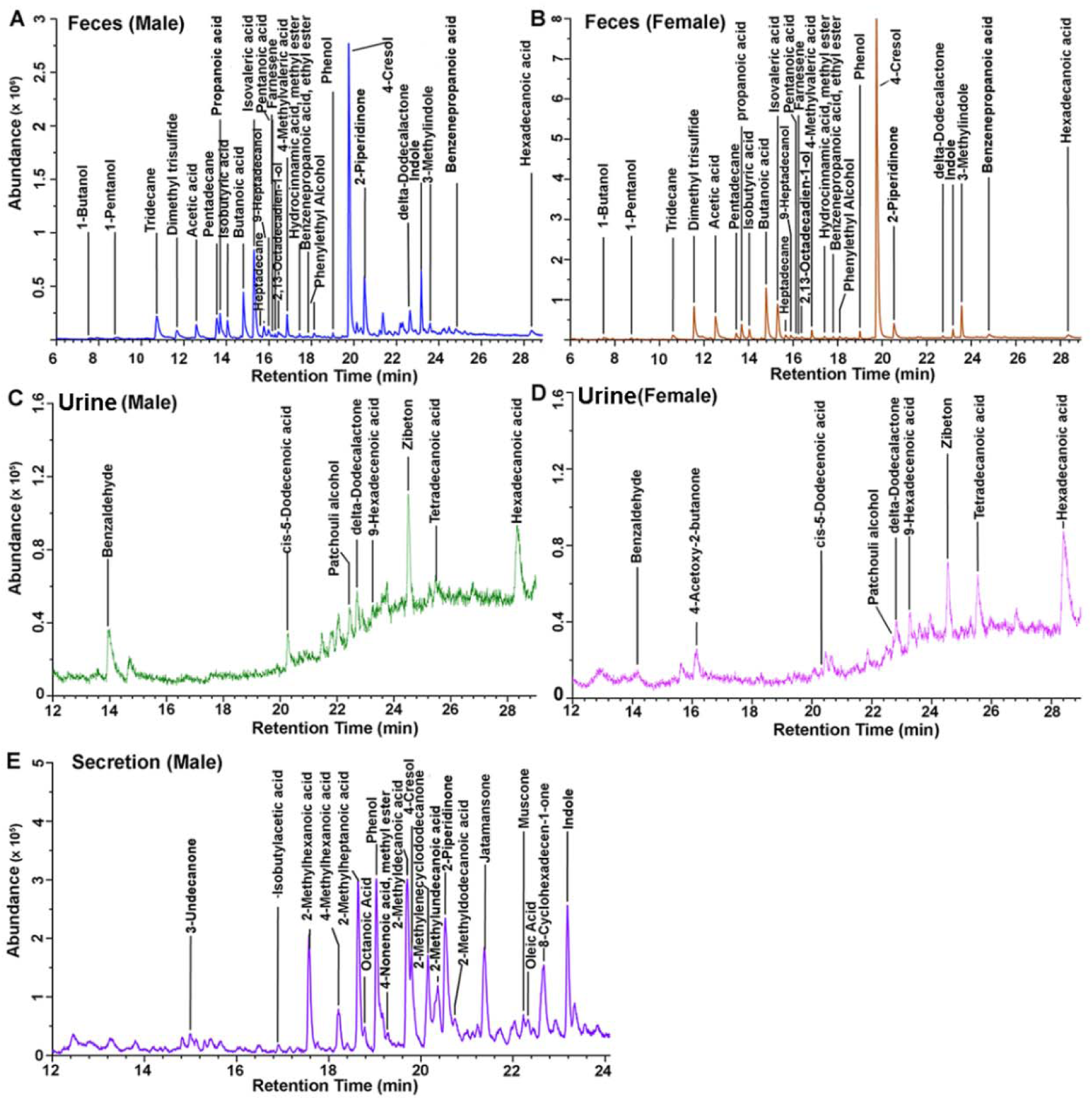
GC-MS analysis. Total ion chromatogram of the feces, urine and anal gland secretion of adult *M. javanica* on a polyethylene glycol column (Table S1).

### Muscone binds to MjavOBP3 with high affinity

A transcriptome analysis identified 34 lipocalin proteins, which belong to 14 different families, including three OBPs, 12 fatty acid-binding proteins, five retinoid-binding proteins, two β-lactoglobulins, and 12 other lipocalin proteins (Table 1). MjavOBP1 was only expressed in the liver, while MjavOBP2 and MjavOBP3 were specifically expressed in the nasal cavity, suggesting their potential olfactory sensing roles (Fig 2A; S2 Table).

**Figure 2.**
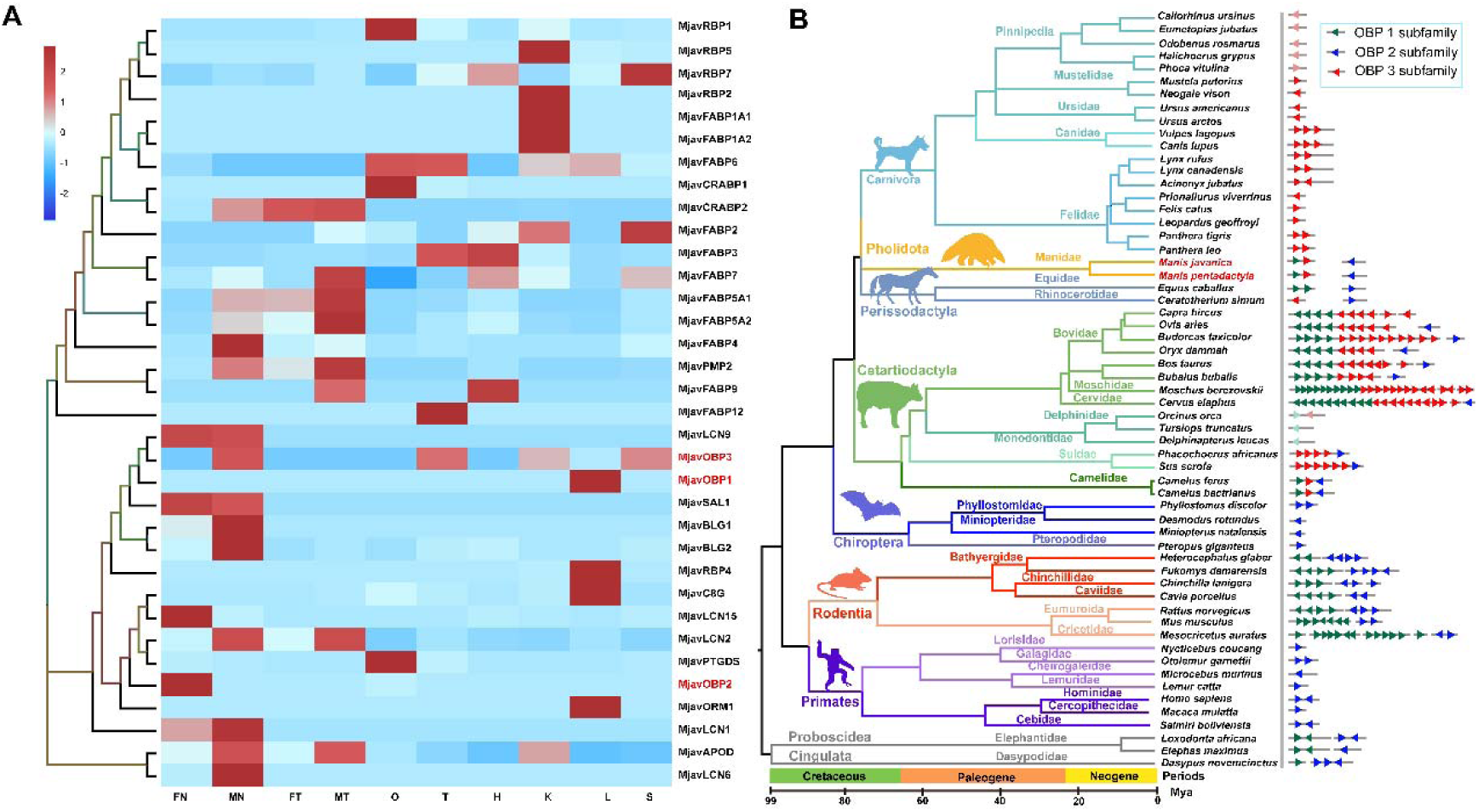
Transcriptome and Homology analysis. (A) Expression profiles of lipocalins in *M.javanica*. (FN: female nasal olfactory epithelium, MN: male nasal olfactory epithelium, FT: female tongue, MT: male tongue, O: ovary, T: testis, H: mixed sample of heart; K: mixed sample of Kidney; L: mixed sample of liver, S: mixed sample of stomach). The left side of the figure is a phylogenetic tree constructed using neighbor-joining for pangolin lipocalins, and the right side is the corresponding gene name (Table S3). (B) Homology analyses of odorant binding proteins (OBPs) in 59 mammalian species. The gene tree constructed for odorant binding proteins using the maximum likelihood method is divided into three clades, corresponding to the OBP1-3 subfamily (Fig. S1, Table S4). The right side of the figure shows the number and location number of different types of OBPs in different color located on the chromosome.

**Table 1.**
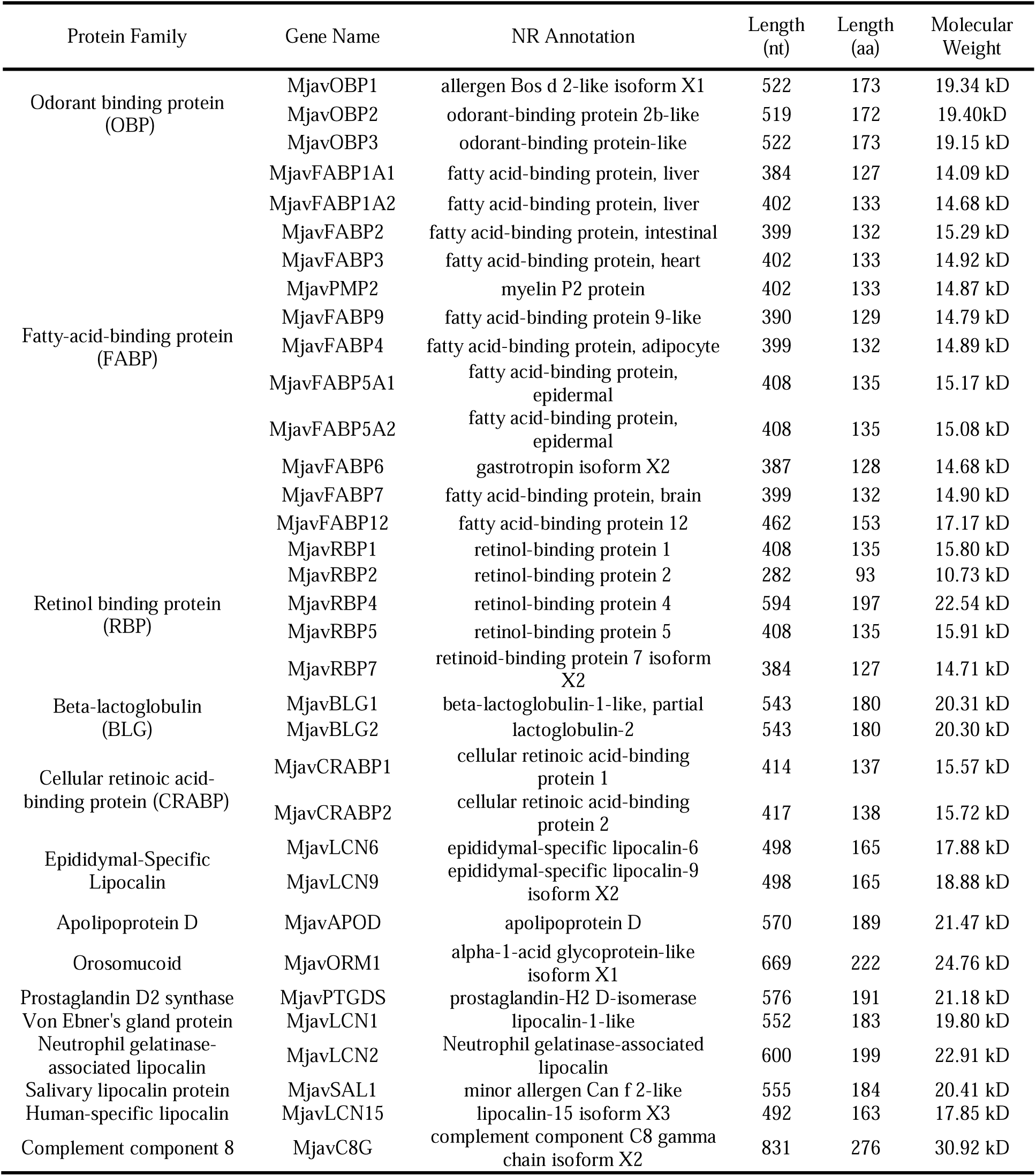
Sunda pangolin lipocalins.

A homology analysis showed that pangolins are most closely related to the order Carnivora, and they diverged during approximately the post-Cretaceous period (74 Mya). The number of OBPs highly varied among different species, ranging from 1 to 27 (S1 Fig; S3 Table). The number of OBPs in Chiroptera, Primates, and Carnivora was close to that of pangolins, and the order Artiodactyla had up to 27 OBPs. In the Cetacea, all the OBPs appeared to be pseudogenes that contained a premature termination codon, suggesting the degeneration of their olfactory-sensing in underwater habits (Fig 2B).

We then analyzed the functional expression of all three MjavOBPs. After purification, sodium dodecyl-sulfate polyacrylamide gel electrophoresis (SDS-PAGE) showed that the MjavOBPs had a target band at approximately 20 kDa (S2 Fig), which was consistent with the predicted length of the amino acid sequence of those proteins. A liquid chromatography-mass spectrometry (LC-MS) analysis showed that the coverage rates of the measured amino acid sequences ranged from 83% to 100% when compared with the reference sequences, validating the MjavOBPs that were purified (Fig 3A-C).

**Figure 3.**
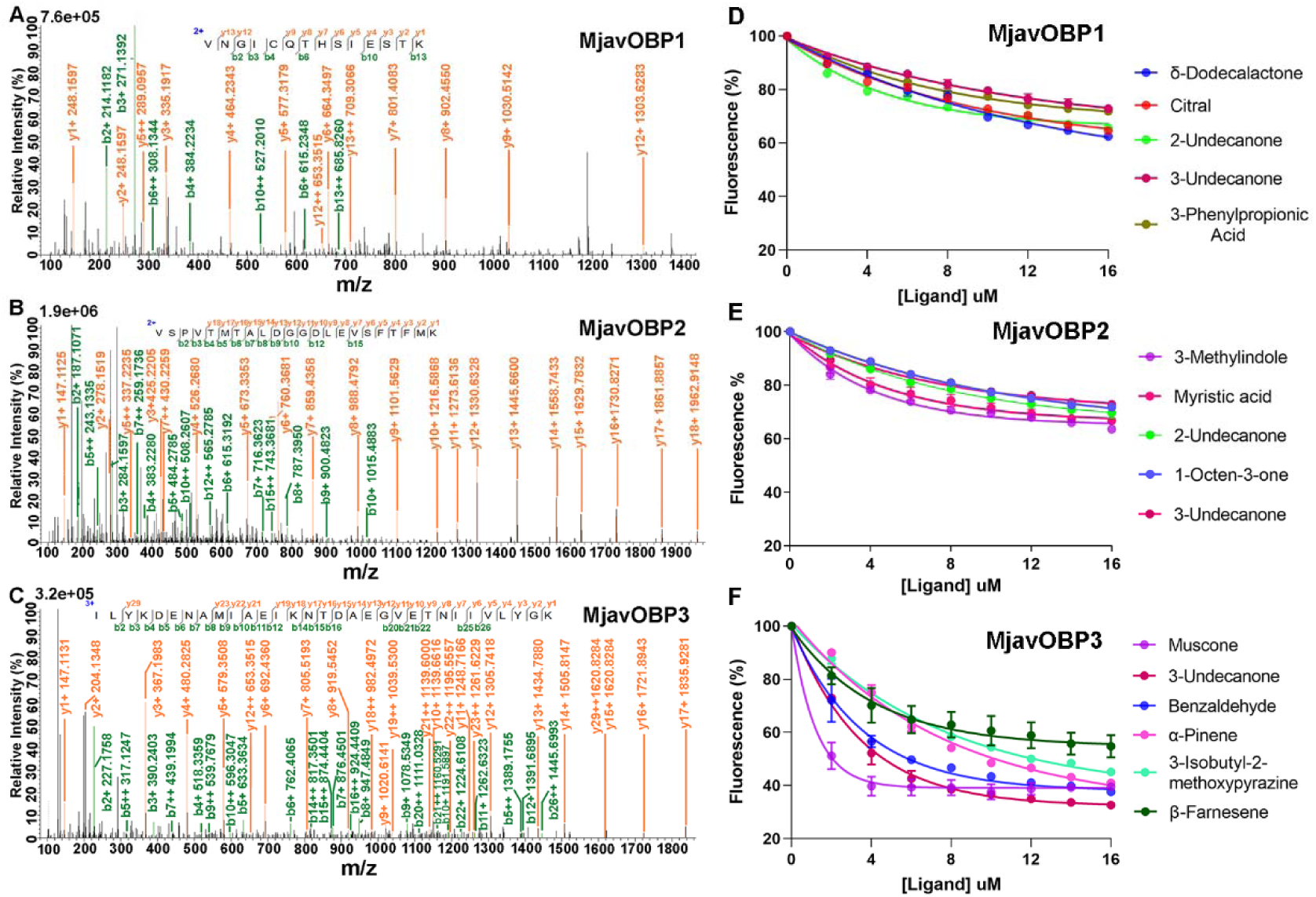
Sequencing analysis and ligand binding ability determination of three MjavOBPs. (A-C) Mass Spectrum of peptides from three MjavOBPs are shown, respectively. The fragments are reported in different color depending on peptides present in *M. javanica* and corresponding b and y ion series. (D-F) Binding curves of all MjavOBPs to different ligands. The ligand names are shown on the right of the curves with different colors, and binding affinity data are listed in Table S6.

Subsequently, three MjavOBPs were screened against a panel of 32 candidate scent-marking and plant volatiles using N-phenyl-1-naphthylamine (1-NPN) as a fluorescent reporter (Table 2, Fig 3D-F; S3 Fig). A competitive binding assay showed that both MjavOBP1 and MjavOBP2 bound to 2-undecanone with K _d_ values of 56.55 ± 4.20 μ M and 54.48 ± 10.75 μ M, respectively. Furthermore, MjavOBP1 had the best binding ability with the urinary volatile δ-dodecalactone (38.74 ± 2.04 μ M) and also bound weakly with other ligands, such as Citral and 1-octen-3-ol. Then, MjavOBP2 had the highest binding affinity for Skatole (40.83 ± 8.93 μ M), and MjavOBP3 showed strong binding (IC50,2.23 ± 0.80 μ M) with muscone, which is an anal gland secretion volatile, and MjavOBP3 also binds to other ketones, sesquiterpenes, and aldehydes, such as 1-octen-3-one, β-farnesene, and Hexanal, with the IC50 value being higher than 2.23 μ M (S4 Table).

**Table 2.**
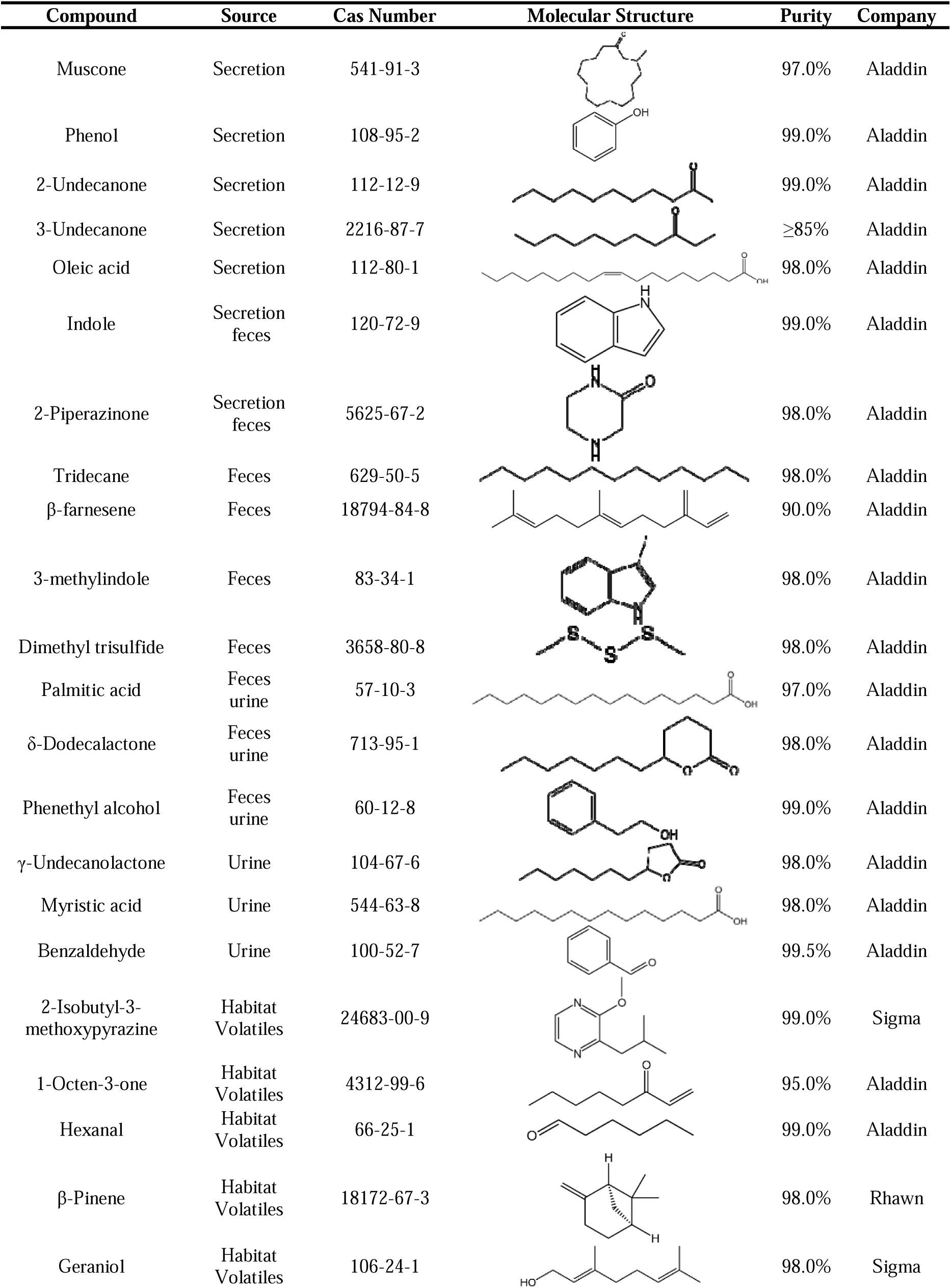
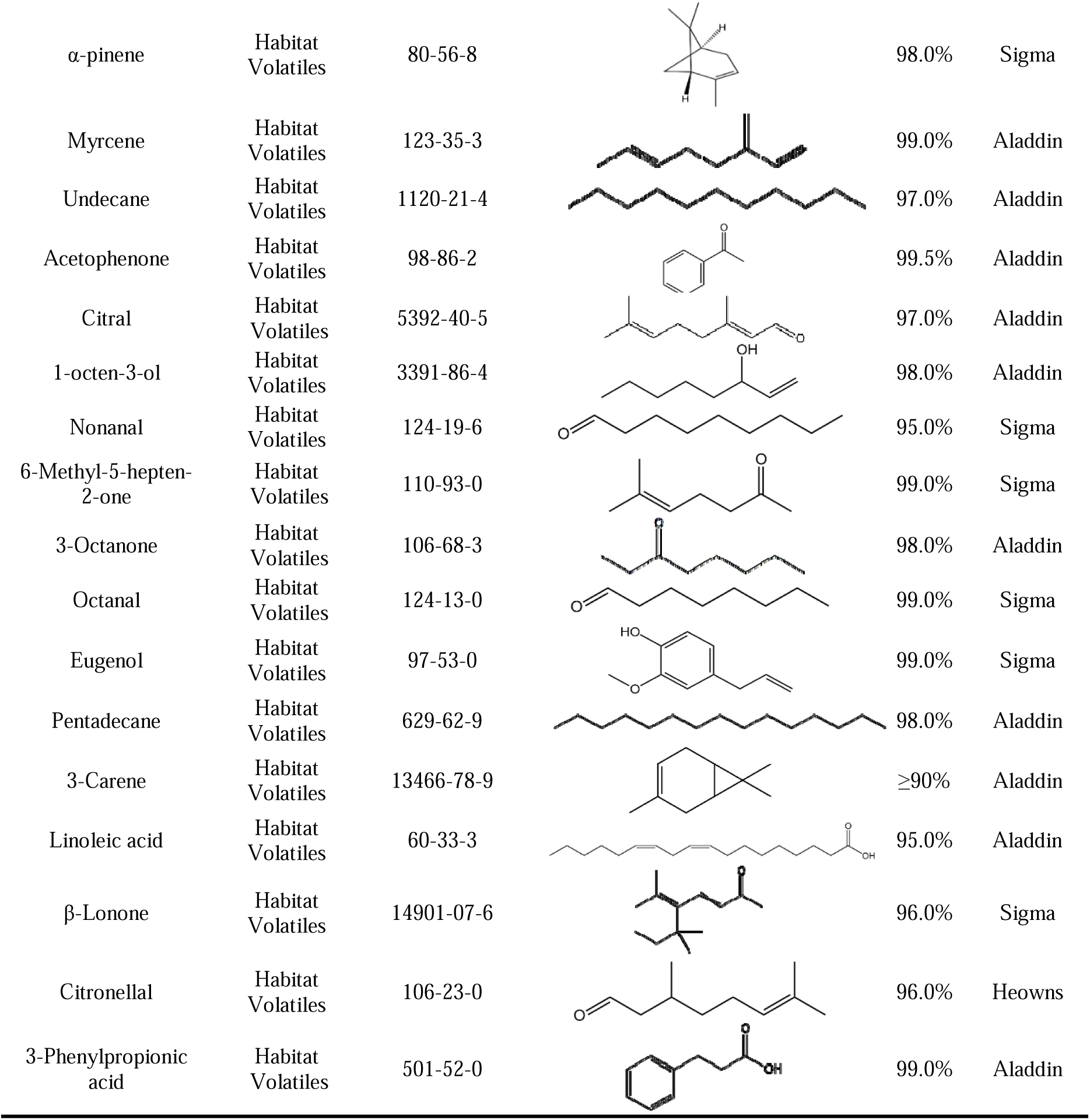
Compounds used for functional validation of odorant binding proteins (OBPs) in Sunda pangolins.

| Compound | Source | Cas Number | Molecular Structure | Purity | Company |
| --- | --- | --- | --- | --- | --- |
| Muscone | Secretion | 541-91-3 |  | 97.0% | Aladdin |
| Phenol | Secretion | 108-95-2 |  | 99.0% | Aladdin |
| 2-Undecanone | Secretion | 112-12-9 |  | 99.0% | Aladdin |
| 3-Undecanone | Secretion | 2216-87-7 |  | ≥85% | Aladdin |
| Oleic acid | Secretion | 112-80-1 |  | 98.0% | Aladdin |
| Indole | Secretion<br>feces | 120-72-9 |  | 99.0% | Aladdin |
| 2-Piperazinone | Secretion<br>feces | 5625-67-2 |  | 98.0% | Aladdin |
| Tridecane | Feces | 629-50-5 |  | 98.0% | Aladdin |
| β-farnesene | Feces | 18794-84-8 |  | 90.0% | Aladdin |
| 3-methylindole | Feces | 83-34-1 |  | 98.0% | Aladdin |
| Dimethyl trisulfide | Feces | 3658-80-8 |  | 98.0% | Aladdin |
| Palmitic acid | Feces<br>urine | 57-10-3 |  | 97.0% | Aladdin |
| δ-Dodecalactone | Feces<br>urine | 713-95-1 |  | 98.0% | Aladdin |
| Phenethyl alcohol | Feces<br>urine | 60-12-8 |  | 99.0% | Aladdin |
| γ-Undecanolactone | Urine | 104-67-6 |  | 98.0% | Aladdin |
| Myristic acid | Urine | 544-63-8 |  | 98.0% | Aladdin |
| Benzaldehyde | Urine | 100-52-7 |  | 99.5% | Aladdin |
| 2-Isobutyl-3-methoxypyrazine | Habitat<br>Volatiles | 24683-00-9 |  | 99.0% | Sigma |
| 1-Octen-3-one | Habitat<br>Volatiles | 4312-99-6 |  | 95.0% | Aladdin |
| Hexanal | Habitat<br>Volatiles | 66-25-1 |  | 99.0% | Aladdin |
| β-Pinene | Habitat<br>Volatiles | 18172-67-3 |  | 98.0% | Rhawn |
| Geraniol | Habitat<br>Volatiles | 106-24-1 |  | 98.0% | Sigma |
| α-pinene | Habitat<br>Volatiles | 80-56-8 | | 98.0% | Sigma |
| Myrcene | Habitat<br>Volatiles | 123-35-3 |  | 99.0% | Aladdin |
| Undecane | Habitat<br>Volatiles | 1120-21-4 |  | 97.0% | Aladdin |
| Acetophenone | Habitat<br>Volatiles | 98-86-2 |  | 99.5% | Aladdin |
| Citral | Habitat<br>Volatiles | 5392-40-5 |  | 97.0% | Aladdin |
| 1-octen-3-ol | Habitat<br>Volatiles | 3391-86-4 |  | 98.0% | Aladdin |
| Nonanal | Habitat<br>Volatiles | 124-19-6 |  | 95.0% | Sigma |
| 6-Methyl-5-hepten-2-one | Habitat<br>Volatiles | 110-93-0 |  | 99.0% | Sigma |
| 3-Octanone | Habitat<br>Volatiles | 106-68-3 |  | 98.0% | Aladdin |
| Octanal | Habitat<br>Volatiles | 124-13-0 |  | 99.0% | Sigma |
| Eugenol | Habitat<br>Volatiles | 97-53-0 |  | 99.0% | Sigma |
| Pentadecane | Habitat<br>Volatiles | 629-62-9 |  | 98.0% | Aladdin |
| 3-Carene | Habitat<br>Volatiles | 13466-78-9 | | $\geq 90\%$ | Aladdin |
| Linoleic acid | Habitat<br>Volatiles | 60-33-3 |  | 95.0% | Aladdin |
| β-Lonone | Habitat<br>Volatiles | 14901-07-6 | | 96.0% | Sigma |
| Citronellal | Habitat<br>Volatiles | 106-23-0 |  | 96.0% | Heowns |
| 3-Phenylpropionic acid | Habitat<br>Volatiles | 501-52-0 |  | 99.0% | Aladdin |

### Behavioral responses of the pangolins to muscone

To examine the behavioral effect of muscone, a trace tracking assay was performed. The results showed that both male and female pangolins visited and stayed in the f5 area more frequently, where was the entrance of the resting place. After placing the control odor source (propylene glycol), both the males and females visited and stayed on the left side of the room more frequently, which was far away from the odor source (b8), suggesting that neither the males nor females were able to recognize the control odor (Fig. 4A and C; S2 and 4 videos). After the muscone was placed, the males could accurately recognize the muscone (area b8, Fig 4 B), and obvious sniffing behavior was observed (S3 Video). In contrast, the females were mainly distributed in f5-6, e5-6, and b5-6 (Fig 4 D), and no preference for muscone was detected (S5 Video).

**Figure 4.**
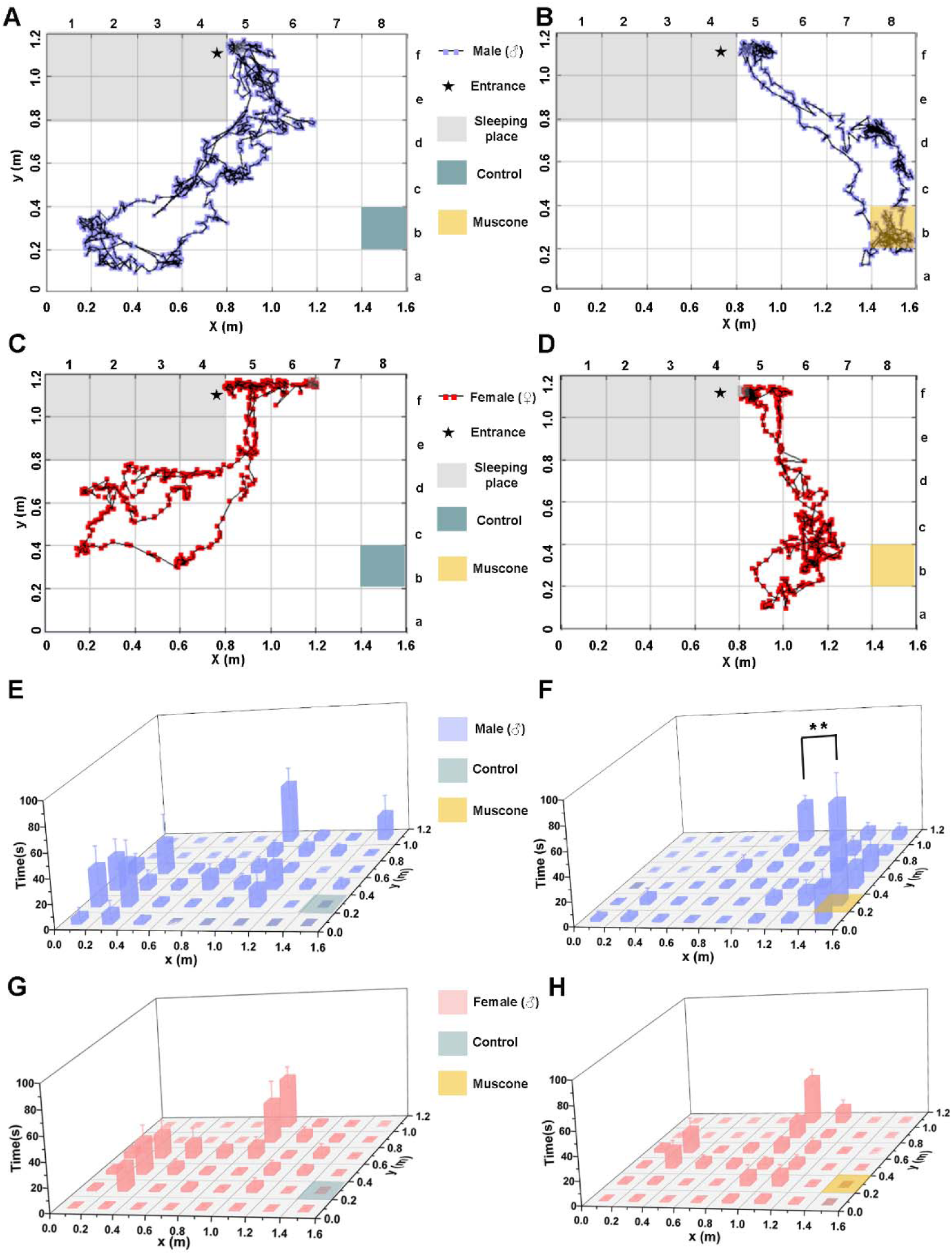
Preferential behavioral responses of female and male pangolins to muscone. (A-D) An individual test pangolin is introduced into the room, can freely enter and exit from the sleeping place (gray area), and muscone odor or control odor (propylene glycol) is placed in b8, and the behavioral trajectories of pangolins to control odors and muscone were recorded separately, ★ represents the entrance, control odor is in cyan, and muscone is in yellow, blue dots represent males, red dots represent females; (E-H) The residence time of pangolins in different areas after placing control odor or muscone. Blue histograms represent males and red histograms represent females. Bars indicate the mean time that a pangolin spent investigating each test sample within a 5-min period, ±S.E.M. Asterisks indicate significant differences between two samples (paired Student’s t-test, **P < 0.01, n=10).

Then, the average staying time of the pangolins in each area was based on three males and three females. In the control group, the highest staying time of the males was in the order of f5 (45.09 ± 12.46 s), b2 (35.62 ± 15.04 s), and b1(31.11 ± 15.23 s), while that for the females was in the order of f5 (38.26 ± 8.25 s), e5 (31.39 ± 15.37 s), and c2 (26.67 ± 10.49 s). The average staying time for both sexes in b8 was 0 (Fig. 4 E and G), suggesting that neither the males nor females could sense the control odor. In the muscone group, the highest staying time for the males was for b8 (78.10 ± 20.04 s), f5 (30.29 ± 5.57 s), and a8 (29.48 ± 9.14 s), and the staying time in the muscone area (b8) was significantly (P < 0.01) higher than that of any of the other areas (Fig. 4 F). In contrast, the highest staying time for the females occurred in f5 (35.72 ± 6.94 s), d2 (18.25 ± 8.85 s), and c2 (15.18 ± 5.78 s), and no staying or sniffing behavior was observed in the b8 area (Fig 4 H; S5 Video). In summary, the behavioral tracking results showed that the male pangolins showed a strong preference for muscone when compared with the control group, and the females could not detect it.

### Key binding sites for muscone on MjavOBP3

The 3D modeling and docking showed that muscone is wrapped in the center of a binding pocket on MjavOBP3 (Fig. 5 A and B; S4 Fig). Additionally, Tyr117 can form a hydrogen bond with muscone, with a molecular distance of 2.8 Å, and the binding of muscone is also attributed to a Pi-alkyl interaction (Phe52), an alkyl interaction (Ile101), and Van der Waal forces (Glu68, Tyr81, and Tyr87; Fig. 5 C). The 100 ns time-evolution root-mean-squared deviation (RMSD) test showed that the MjavOBP3-muscone complex systems experienced a large fluctuation during 0–14 ns and reached equilibrium at 2.26 Å within 50 ns. The standard deviation of the RMSD after 50 ns was 0.14 Å, indicating the stability and reliability of the MD trajectory (Fig. 5 D). In the equilibrium phase, Tyr117 formed a hydrogen bond, and the other non-polar residues (Phe52, Glu68, Tyr81, Tyr87, and Ile101) formed Van der Waal forces and a (pi)-alkyl bond, which also contributed to the MjavOBP3-muscone interactions (Fig. 5 E). The binding free energy of the MjavOBP3-muscone complex was predicted to be -39.42 kcal/mol (S5 Table). Subsequently, the residues with a total energy contribution (ΔG bind) exceeding −1.00 kcal/mol were chosen for further validation (Fig. 5 F), which included Tyr117, Phe52, Ile101, Glu68, Tyr81, Tyr87, and two negative control residues (Met97, inside the binding pocket; Lys130, far away from the binding pocket).

**Figure 5.**
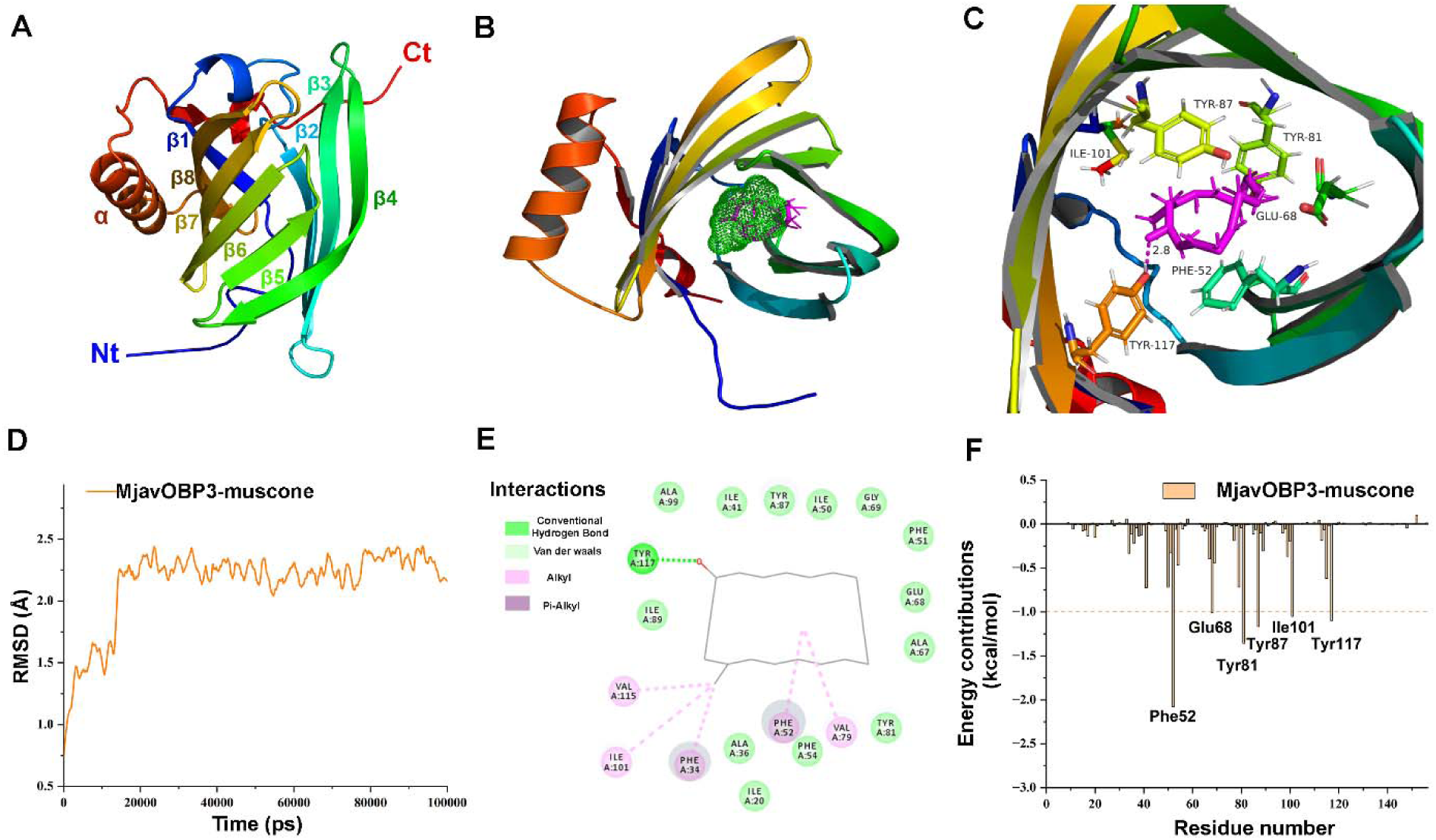
Prediction of key sites in the MjavOBP3-Muccone complex. (A) The 3D structure of MjavOBP3 presents a barrel-like structure, mainly composed of 8 β-sheets (β1-β8) and 1 α-helix; (B) The predicted binding pocket is shown in green. The ligand Muscone (dark pink) is tightly wrapped by the binding pocket; (C) 3D visual analysis of the binding mode of MjavOBP3 and muscone. Important amino acid residues and muscone are shown as baseballs, in which hydrogen bonds are represented by pink dotted lines, and molecular distances are calculated; (D) The root-mean-square deviation (RMSD) curve of the MjavOBP3−Muscon complex. (E) Protein-ligand interaction diagram of MjavOBP3 bound to muscone at steady state; (F) Total energy contribution of each amino acid residue in the MjavOBP3−muscone complexes. The residues with a total energy contribution of more than −1.00 kcal/mol are marked.

When compared with the wild type, the binding affinity of muscone in the four mutations (Y117F, F52A, Y87A, and Y81A) was significantly decreased (P < 0.01; Fig. 6 A-C; Fig. S5), which shows that Y117F does not bind to muscone with a K _d_ value of 116.78 ± 4.41 μ M. The fluorescence binding results were highly consistent with the modeling and docking predictions, indicating that the Tyr117, Phe52, Tyr87, and Tyr81 residues are crucial for the binding of muscone, and the Tyr117 residues contributed the most to the binding process.

**Figure 6.**
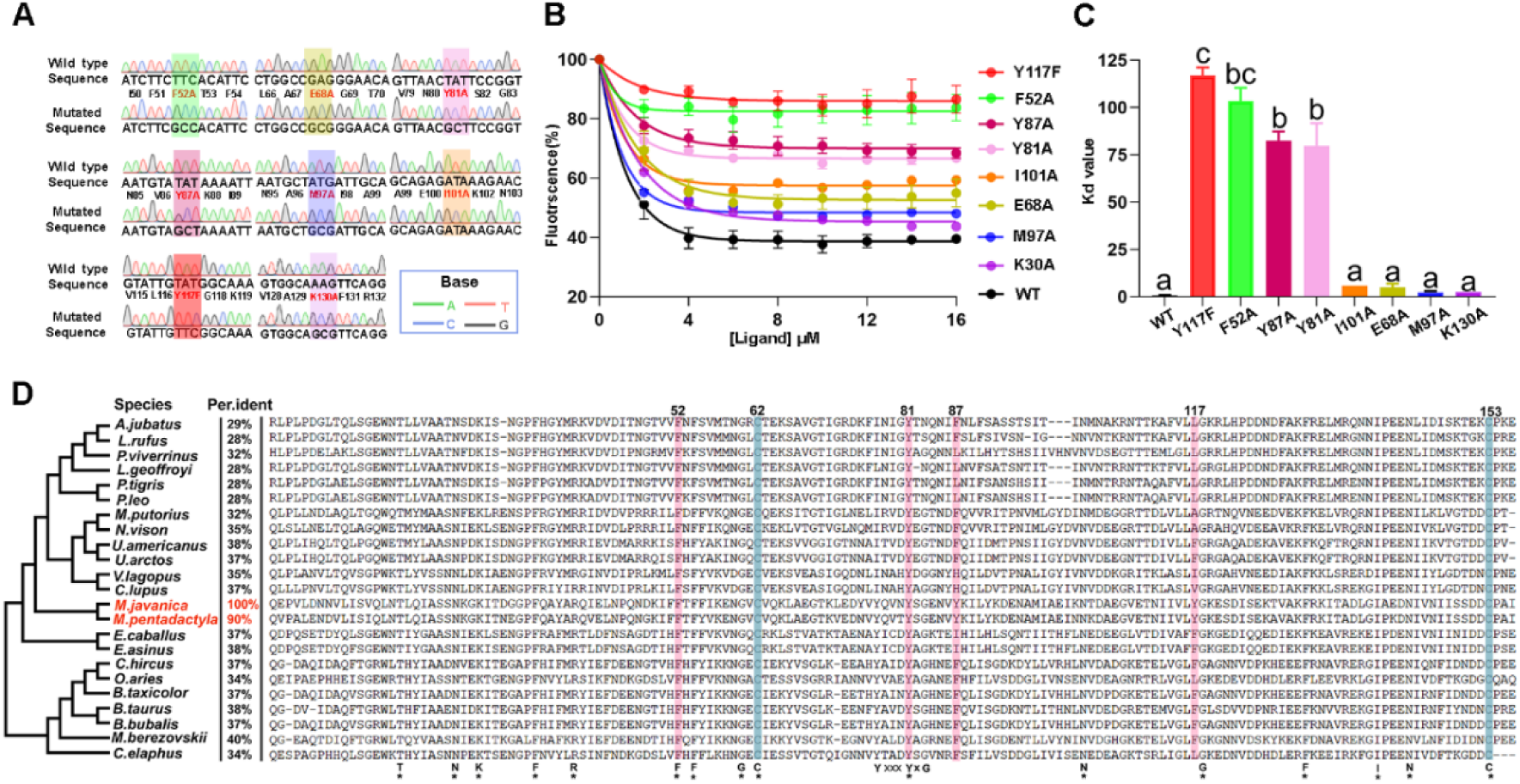
Verification of key sites of MjavOBP3-muscone complex. (A) Target site in the MjavOBP3 gene (wild type) and results obtained from the mutagenesis of MjavOBP3 through site-directed mutagenesis, and bases A, T, C and G are represented by green, pink, blue and black respectively, and the eight mutation positions are represented by different color; (B) Binding curves of MjavOBP3 mutants to muscone. The lines (different colors) represent the different mutants, and the line (black) represents the wild type (WT); (C) Comparison of the binding affinities, which were indicated by K d value, of MjavOBP3 WT and mutants to muscone, respectively. The significant differences were compared using one-way analysis of variance (ANOVA) with Tukey’s honestly test and indicated by different lowercase letters (p-value < 0.01, n =3); (D) Comparison of the amino acid sequence similarity of MjavOBP3 homologous proteins in Laurasia. The sequence similarity is expressed by per ident. The stars (*) show the same amino acids in all proteins. Conserved cysteine amino acid sites involved in the formation of disulfide bridges are shown in light blue, while Key binding sites in MjavOBP3 for Muscone are shown in light pink

The amino acids alignment on MjavOBP3 showed a conserved YxxxYxG motif (positions 77–83) and 16 conserved residues (indicated by *) among its orthologues (Fig. 6 D). The similarity of the homologous OBP3s among the different species ranged from 28% to 90%, and the highest similarity was with its close relative, *Manis pentadactyla*. The above four key residues (Tyr117, Phe52, Tyr87, and Tyr81) were highly conserved between *M. javanica* and *M. pentadactyla*, suggesting a similar olfactory sensing pathway for muscone. Among the key binding residues, Phe52 (Pi-alkyl) and Tyr81 (van der Waals force) were highly conserved among the mammals, indicating their crucial role in the OBP3 orthologues. In contrast, Tyr117 (hydrogen bond) was only conserved in *M. javanica* and *M. pentadactyla*, suggesting that it contributes the most to the muscone binding and has a crucial role in specific sensing of muscone in pangolins.

## Discussion

The volatile organic compounds (VOCs) that were identified from the urine, feces, and anal gland secretions of the pangolins were mainly carboxylic acids, ketones, and alcohols, which were followed by lactones and aldehydes. Among them, carboxylic acids, aldehydes, phenols, and indoles are often detected in mammal excretions [26] and these VOCs may not convey intraspecies information. Notably, some of the VOCs, such as δ-dodecalactone, farnesene, and Muscone, are rare in nature or pangolin-specific, indicating their potential pheromone role. Specifically, δ-dodecalactone may act as a scent-marking signal in male Bengal tigers (*Panthera tigris*) [27], and Farnesene, which is derived from the urine of male mice, was shown to be a territory-marking pheromone [28]. Muscone is a rare VOC in mammal secretions, and only musk deer have been recorded to synthesize it using long-chain fatty acids [29]; they use it for territory marking and female attraction [30]. The food sources of pangolins (ants and termites) do not contain any muscone-relevant compounds [31, 32], indicating that there might be a specific biosynthesis pathway for muscone in pangolins, which would reflect their unique biological function.

The OBPs’ main function is to increase the solubility of odorants and transport odorants to olfactory receptors in a hydrophilic environment [23]. The number of OBPs varies greatly among the different animal groups; for instance, the number of OBPs in insects mainly ranges from 40 to 60 [33], while the number of OBPs in mammals is comparatively small. Our study showed that the number of OBPs in the orders Pholidota, Carnivora, Perissodactyla, Chiroptera, and Primates were only 1-3, and the reduction of OBPs in these species was likely due to an insufficient olfactory transport ability to varied environmental volatiles, which limited their function to pheromone carrying [23]. Previous studies have shown that OBPs in mammals are not synthesized in the olfactory organ but are synthesized in the vomeronasal organ (VON) and nearby areas [34–37]. The VON is the main pheromone-sensing organ in mammals, and it shortens the transportation distance of the pheromone. A recent study showed that AimelOBP3 in giant pandas is bound with linear aldehydes, and they may be involved in pheromone recognition [22]. Pangolins are fossorial, and their vision quality is low but their olfactory system is well developed, and, thus, chemical signals play a key role in their intraspecies communications [20]. The fluorescence binding assay showed that MjavOBP3 had the highest binding affinity to muscone, suggesting that muscone has a putative pheromone-sensing role.

To further explore the behavioral effect of muscone on pangolins, a well-controlled behavioral assay was performed on this endangered species. The behavioral tracking assay indicated that male pangolins could recognize muscone but females could not, which was consistent with the fact that MjavOBP3 is only expressed in the olfactory epithelium of males. Muscone is released by the male anal glands, and the anal gland is a common organ in mammals, its main function is to synthesize anal gland secretions and lubricate the feces. However, in some cases, the anal gland is also responsible for territory marking and intraspecies recognition (such as in *Meles meles* and *Herpestes javanicus*) [38, 39]. Male pangolins prefer to live in solitary, and they usually have a fixed living place with a home range of approximately 40 ha [21]. Field observations have shown that pangolins are territorial animals, and males sometimes fight with each other for territory. Considering that pangolins have degraded vision but a well-developed olfaction, thus, territorial marking is believed to rely on chemical signals. Similar behavior was also observed in other Carnivora mammals, and they are close relatives to Pholidota. For example, *Crocuta crocuta* and *Canis lupus* use anal gland secretion for scent and territory marking [40, 41]. The fluoresce binding assays showed that muscone is the best ligand for MjavOBP3, and it is a macrocyclic ketone that is composed of 15 carbons with a slow release and long-lasting aroma, which is consistent with the persistence characteristics of scent-marking pheromones [42]. Moreover, muscone is a rare chemical in nature and has only been detected in a few mammals, such as muskrats (*Ondatra zibethicus*) and musk deer, where it is used for scent marking [29]. Therefore, we hypothesized that muscone is a male scent-marking pheromone in *M. javanica*.

Reverse chemical ecology approaches were applied to identify the pheromone in pangolins. The current conservation strategy would benefit from understanding how male pangolins communicate with each other via chemical signals at a molecular level. The discovery of scent-marking pheromones in pangolins could be used to monitor this endangered species [43, 44], and a scent-based pheromone trap could be developed for the observation and relocation of this endangered animal.

## Materials and Methods

### Ethics statement

All the animal procedures were approved by the ethics committee for animal experiments at the Terrestrial Wildlife Rescue and Epidemic Diseases Surveillance Center of Guangxi, and the recommended guidelines were followed. (S6 Fig)

### Gas chromatography-mass spectrometry analysis

Feces, urine, and anal secretions from Sunda pangolins were collected in January 2021. The individuals that were used in this study were three adult males and three adult females, and the detailed information for each individual is listed in S6 Table. The volatile samples were collected in a 10 mL bottle and adsorbed using a solid-phase microextraction syringe (Supelco 50/30 μm DVB/CAR/PDMS, Manual Holder, 3pk, USA) for 30 min. The samples were analyzed using a GC-MS chromatograph (Agilent 6890N/5973I, USA), and the temperature was maintained isothermally at 80 °C for 3 min and increased to 350 °C at a rate of 10 °C/min for 27 min. The retention time of each peak was used to identify the relevant chemicals based on the National Institute of Standards and Technology database (http://webbook.nist.gov/chemistry), and the relative peak areas were used to assess the abundance of each volatile.

### Transcriptome analysis

Frozen samples of one male (already dead when intercepted by customs in November 2020) and one female (unable to eat due to intestinal disease and died in December 2020) were used to extract the total RNA. Nine tissues from the frozen samples mentioned above were used for RNA sequencing, including the nasal olfactory epithelium of the male and female, tongue of the male and female, ovary, testis, and sex-mixed samples of the brain, heart, kidney, liver, and stomach. The total RNA was extracted using TRIzol reagent (Invitrogen, USA), and the complementary DNA libraries were constructed using magnetic beads and Illumina’s NEBNext Ultra RNA Library Prep Kit (Illumina, USA). The RNA sequencing was conducted on an Illumina platform, and the genome sequence of *M. javanica* (GCF_014570535.1) was used as a reference for annotations. Then, Fragments Per Kilobase of transcript per Million mapped reads (FPKM) was used to calculate the gene expression level.

The homology analysis was performed on an OBP family among 59 mammalian species. Mafft V7 (European Bioinformatics Institute, UK) was used to compare the amino acid sequences, and TBtools-II (Chengjie Chen) was used to construct the maximum likelihood tree with 1000 bootstrap runs and default settings. Meanwhile, the Time Tree of Life database (https://timetree.org/) was used to construct the species tree.

### Gene cloning, expression, and purification

The open reading frame of MjavOBP1-3 was obtained by polymerase chain reaction (PCR) with specific primers that were designed by Primer Premier 5.0 (PREMIER Biosoft International, USA; Table S9) with 2 × TransTaq HiFi PCR SuperMix I (Transgen Bio, China), and they were subcloned to pET30a (Novagen, Germany) between EcoRI and XhoI (NEB, USA) using a pEASY-Basic Seamless Cloning and Assembly Kit (Transgen Bio, China). The plasmids containing the target OBPs were subsequently transformed into BL21(DE3) competent cells and induced by 1 mM isopropyl β-D-1-thiogalactopyranoside at 34 °C for 10 h. The protein expression was checked using 15% SDS-PAGE and loaded onto a Ni-TED 6FF pre-packed chromatography column (Sangon Bio, China) in a High-Performance Liquid Chromatography system (ӒKTA pure, Sweden). The His tag was removed by Recombinant Bovine Enterokinase (Sangon Bio, China).

### Liquid chromatography-mass spectrometry analysis

The LC-MS was conducted according to Bortolussi et al. with a few modifications [45]. In brief, the gel was destained, washed, alkylated, and trypsinized. The samples were then desalted with a Zip-TipC18 column (Sigma Aldrich, USA) and eluted with a 67% acetonitrile solution containing 2% formic acid. The protein sample was analyzed by a nano LC-ESI-Q Orbitrap tandem mass spectrometry (MS/MS) platform that consisted of the Dionex Ultimate 3000 HPLC RSLC nano system (Thermo Fisher Scientific, USA) and Q-Exactive Plus mass spectrometer (Thermo Fisher Scientific) mounted to a Nanoflex ion source (Thermo Fisher Scientific). The sample was first bound to a C18 Reversed Phase Column (Acclaim PepMap RSLC, 75 μ M × 150 mm, 2 μ M 100 Å; Thermo Fisher Scientific) and eluted by gradient acetonitrile (from 0% to 80% over 80 min) containing 0.1% formic acid at 1 mL/min. The mass spectrometer was operated under the data-dependent analysis mode (MS1 scan resolution: 70000; scanning range: 350–2000 m/z), which was followed by an MS/MS scan of the ten most abundant ions in high-energy collisional dissociation mode (maximum ion injection time: 50 ms; collision energy: 28 eV; dynamic exclusion: 25 s). Finally, the mass spectrum was converted into mgf format using ProteoWizard 3.0 software [46] and imported into Mascot v. 2.6.1 (Matrix Science, UK) for protein identification. The settings were as follows: database, uniprot; enzyme, trypsin; maximal number of missed cleavages, 1; MS tolerance, 20 ppm; MS/MS tolerance, 0.05 Da; protein score, ≥ 95%).

### Fluorescence binding assays

The reaction was conducted in a 2 mL transparent quartz cell (Hellma, Germany). A mixture was constructed of 2 μ M protein dissolved in 50 mM Tris·HCl (pH 7.4) and 2 μ M probe 1-NPN dissolved in 50 mM Tris·HCl (pH 7.4), ligands were dissolved in methanol eight times to ensure the final content of the ligands was 2-16 μ M under a Fluoromax-4 spectrofluorometer (HORIBA Jobin Yvon, USA), and they were added to this mixture. Then, 1-NPN was used as a fluorescent reporter. The excitation wavelength was set to 337 nm, and the emission spectrum was recorded between 380 and 450 nm under an emission slit of 5 nm. The dissociation constants of the competitors were calculated using the equation K _d_ = IC_50_ / (1+ [1-NPN]/K_1-NPN_), where IC_50_ is the concentration of ligands halving the initial fluorescence value of 1-NPN, [1-NPN] is the free concentration of 1-NPN, and K_1-NPN_ is the dissociation constant of the complex protein/1-NPN. GraphPad Prism 9.0 software (GraphPad by Dotmatics, USA) was used for the K _d_ calculation.

### Behavioral tracking

The behavioral tracking assay was conducted on six individuals (S6 Table). The experiment was conducted during the activity peak of the pangolins (8 pm to 2 am) in a 1.6 × 1.2 m breeding room. The room was artificially divided into 48 small regions (20 × 20 cm) to record the visiting time of the different areas. Then, 450 μg of muscone dissolved in 40 μL propylene glycol was used as the odor stimulus, and pure propylene glycol was used as a negative control. The stimulus was added to an ethanol-cleaned plastic round box (2.4 cm in diameter, 1.2 cm in height) and placed at the corner of the room, and the behavior of the pangolins was recorded as soon as they left the resting area. A two-day interval was used between each experiment to avoid conditional reflex effects. The behaviors were recorded using an infrared camera (Cuddeback Expert, USA) under dark conditions, with at least three technical replicates for the six individuals mentioned above. After recording, the behavioral path of the pangolins was tracked by the position of their noses with Tracker 6.1.5 software (https://physlets.org/tracker/trackerJS/; supplementary video 1). The visiting frequency was calculated using the average staying time in each region.

### Molecular modeling and docking of MjavOBP3

The three-dimensional structure of MjavOBP3 was predicted using the Alphafold2 platform based on the amino acid sequence, and the highest score model was selected from the top five. The topology of the protein model was optimized by Amber 20 [47], and the stereochemical quality of the modeled protein structure was verified by the PROCHECK method. Next, Sybyl v7.3 (Tripos, USA) was applied to calculate the interaction between the protein and ligand; the muscone was energy minimized, then it was docked to MjavOBP3 with “auto” binding cavity settings, and the total score was used to evaluate the binding affinity of the ligand to the target protein. Finally, the modeling and docking results were visualized by PyMOL (DeLano Scientific LLC).

### Molecular dynamic simulation and site-directed mutagenesis

The topology fields of the ligand and protein were generated under the AMBER ff14SB force field and general AMBER force field in Amber 20 software, respectively, and then the complex was immersed in a cubic box with TIP3P water molecules. Then, Na^+^ was added to neutralize the entire system, and the energy of the complex was minimized by the steepest descent and conjugate gradient methods. Next, the system was heated from 0 to 300 K and the MD simulations were performed at constant 300 K and 1.0 Atmos for 100 ns using the PMEMD module. The RMSD of MjavOBP3-muscone was calculated to assess the equilibrium of the MD trajectories. The binding free energy between the residues and muscone was calculated using the Molecular Mechanics Poisson-Boltzmann Surface Area method [48]. The hydrogen bonding sites and the residues that contributed more than -1.00 kcal/mol to the binding free energy were selected for the mutagenesis validation. Site-directed mutagenesis was achieved by using the Fast Mutagenesis System Kit (TransGen, China), and the mutagenesis primers were designed by Primer Premier 5.0 (S7 Table).

## Acknowledgments

We thank all the reviewers for comments on the manuscript and Yuxin Zhou, Xiaoyan Zhu and Haoqin Ke for their help with molecular biology experiments. as well as MJEditor (www.mjeditor.com) for providing English editing services during the preparation of this manuscript.

## Supporting information

**S1 Fig. Mammalian OBP gene tree based on Maximal likelihood method**

**S2 Fig. SDS/PAGE of MjavOBP1-3 before and after purification**

**S3 Fig. Binding curve of 1-NPN to all MjavOBP1-3**

**S4 Fig. Ramachandran plot of MjavOBP3 model**

**S5 Fig. Binding curve of 1-NPN to MjavOBP3 WT and Mutants**

**S6 Fig. Ethics statement**

**S1 Table (Microsoft Excel format). Determination of VOCs in urine, feces and anal gland secretions in pangolins.**

**S2 Table. (Microsoft Excel format). The expression levels of lipocalins in Sunda pangolins from different tissues.**

**S3 Table. (Microsoft Excel format). List of identified mammalian OBP genes.**

**S4 Table. (Microsoft Excel format). [IC]50 and Kd values between three MjavOBP1-3 and candidate ligands.**

**S5 Table. (Microsoft Excel format). Binding free energy of MjavOBP3-muscone complex.**

**S6 Table. (Microsoft Excel format). Individual information used for experimental pangolins.**

**S7 Table. (Microsoft Excel format). Primers used for plasmid construction of MjavOBP1-3 and mutants.**

**S1 Video (separate files). Pangolin activity videos converted into tracks.**

**S2 Video (separate files). Video of male pangolin’s behavioral response to control odor.**

**S3 Video (separate files). Video of male pangolin’s behavioral response to muscone.**

**S4 Video (separate files). Video of female pangolin’s behavioral response to control odor.**

**S5 Video5 (separate files). Video of female pangolin’s behavioral response to muscone.**

