## supplementary material1 for "Muscone-specific olfactory protein reveals the putative scent-marking pheromone in the Sunda pangolin (*Manis javanica*)"

**This PDF file includes:**

S1 Fig. Mammalian OBP gene tree based on Maximal likelihood method

S2 Fig. SDS/PAGE of MjavOBP1-3 before and after purification

S3 Fig. Binding curve of 1-NPN to all MjavOBP1-3

S4 Fig. Ramachandran plot of MjavOBP3 model

S5 Fig. Binding curve of 1-NPN to MjavOBP3 WT and mutants.

S6 Fig. Ethics statement

**Other supporting materials for this manuscript include the following:**

S1 Table (Microsoft Excel format). Determination of VOCs in urine, feces and anal gland secretions in pangolins.

S2 Table. (Microsoft Excel format). The expression levels of lipocalins in Sunda pangolins from different tissues.

S3 Table. (Microsoft Excel format). List of identified mammalian OBP genes.

S4 Table. (Microsoft Excel format). [IC]50 and K _d_ values between three MjavOBP1-3 and candidate ligands.

S5 Table. (Microsoft Excel format). Binding free energy of MjavOBP3-muscone complex.

S6 Table. (Microsoft Excel format). Individual information used for experimental

pangolins.

S7 Table. (Microsoft Excel format). Primers used for plasmid construction of MjavOBP1-3 and mutants.

S1 Video. (separate files). Pangolin activity videos converted into tracks.

S2 Video. (separate files). Video of male pangolin's behavioral response to control odor.

S3 Video. (separate files). Video of male pangolin's behavioral response to muscone.

S4 Video (separate files). Video of female pangolin's behavioral response to control odor.

S5 Video (separate files). Video of female pangolin's behavioral response to muscone.

**Supporting Information Text**
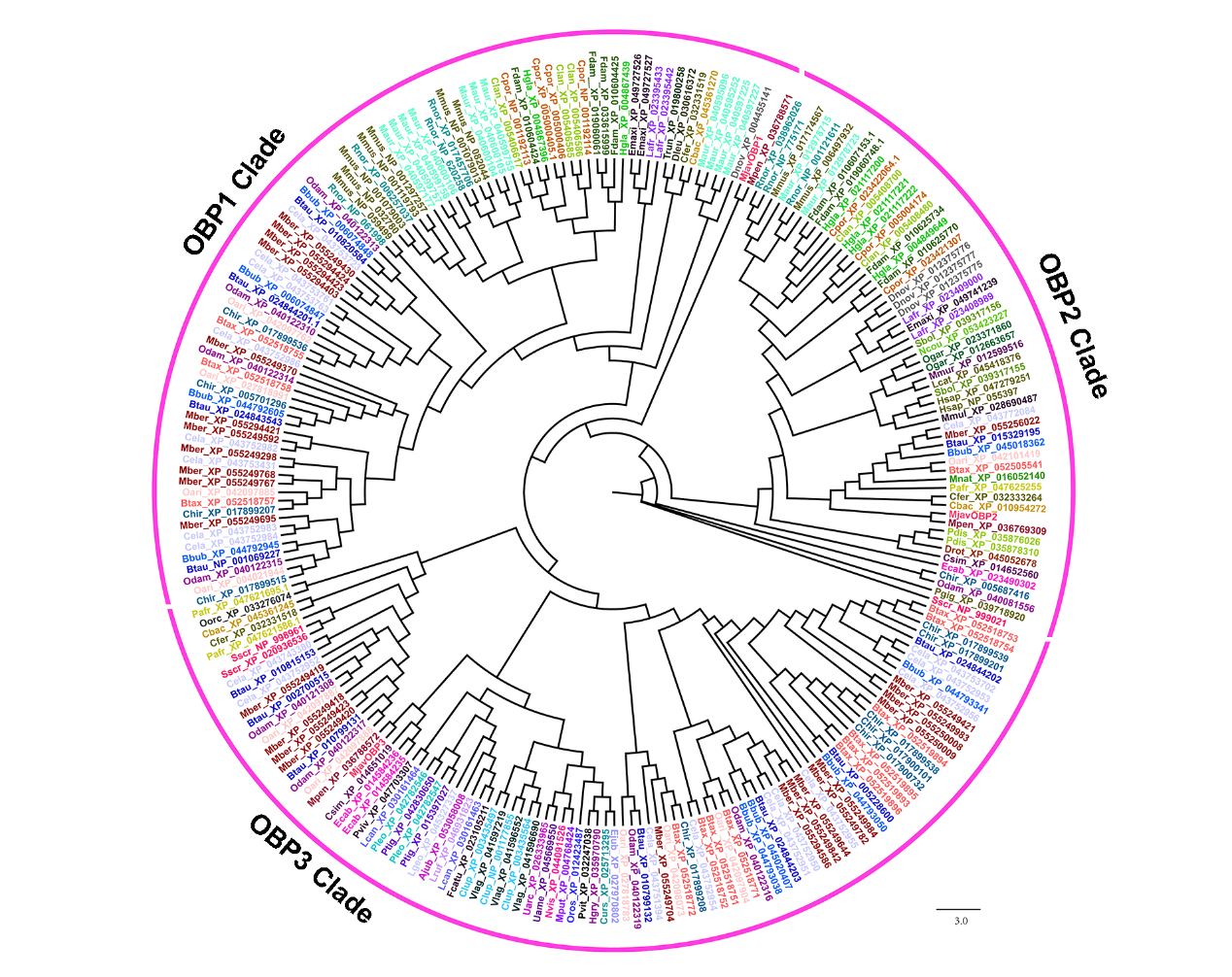

Fig. S1. Mammalian OBP gene tree based on Maximal likelihood method. The phylogenetic tree of MjavOBP1-3 (red) and other mammalian odorant binding proteins odorant-binding proteins (OBPs) (different colors). The tree was constructed using the Maximal likelihood method. The bootstrap repeats 1,000 times. GenBank accession numbers of OBPs used are listed in Table S3.

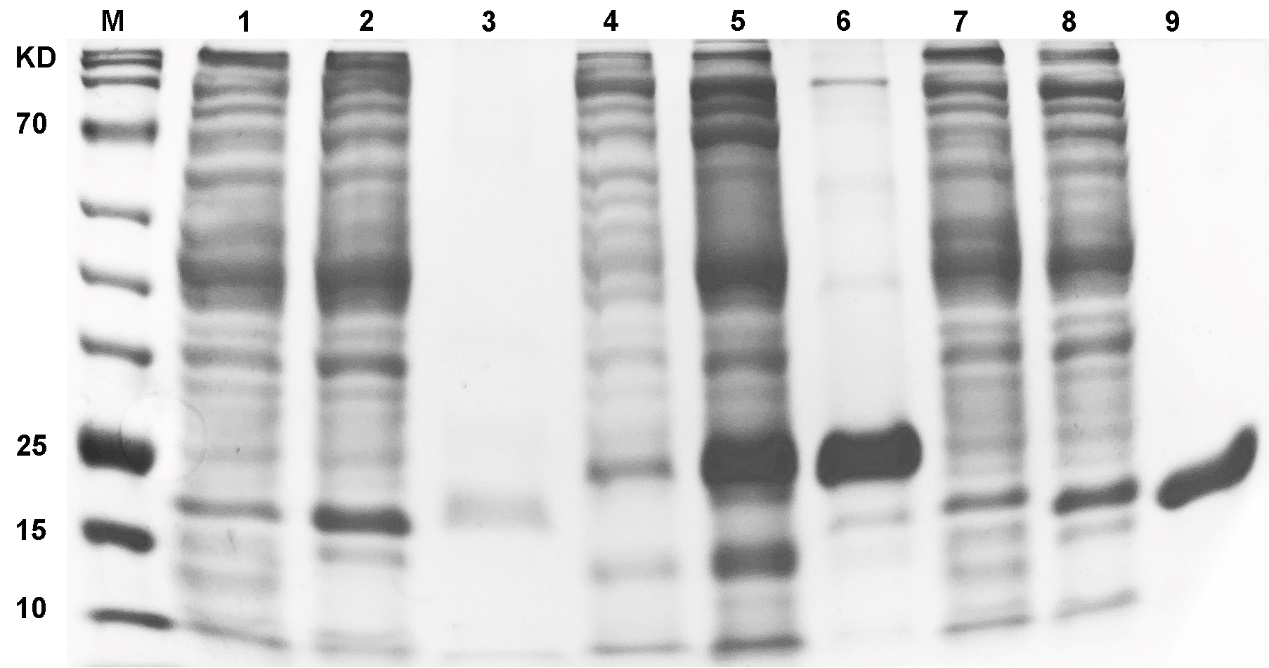

Fig. S2. SDS/PAGE of MjavOBP1-3 before and after purification. Expression and purification of all MjavOBPs. M: 180 kD prestained protein marker;1, 4 and 7 represent the bands of MjavOBP1-3 before induction with IPTG, respectively; 2, 5 and 8 represent the bands of MjavOBP1-3 after induction with IPTG, respectively; 3, 6 and 9 represent the bands of MjavOBP1-3 after purification.

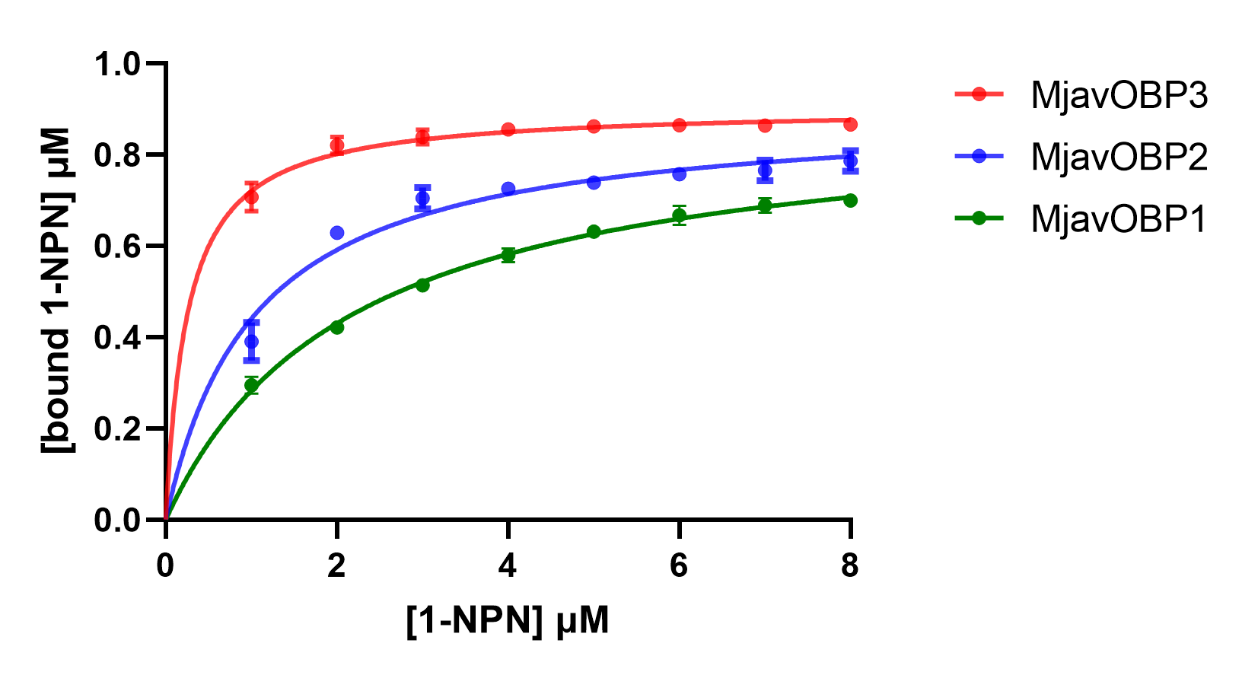

Fig. S3. Binding curve of 1-NPN to all MjavOBPs. The dissociation constants (K_d_) were 2.18±0.25 μM for MjavOBP1, 1.04±0.26 μM for MjavOBP2, 0.42±0.09 μM for MjavOBP3.

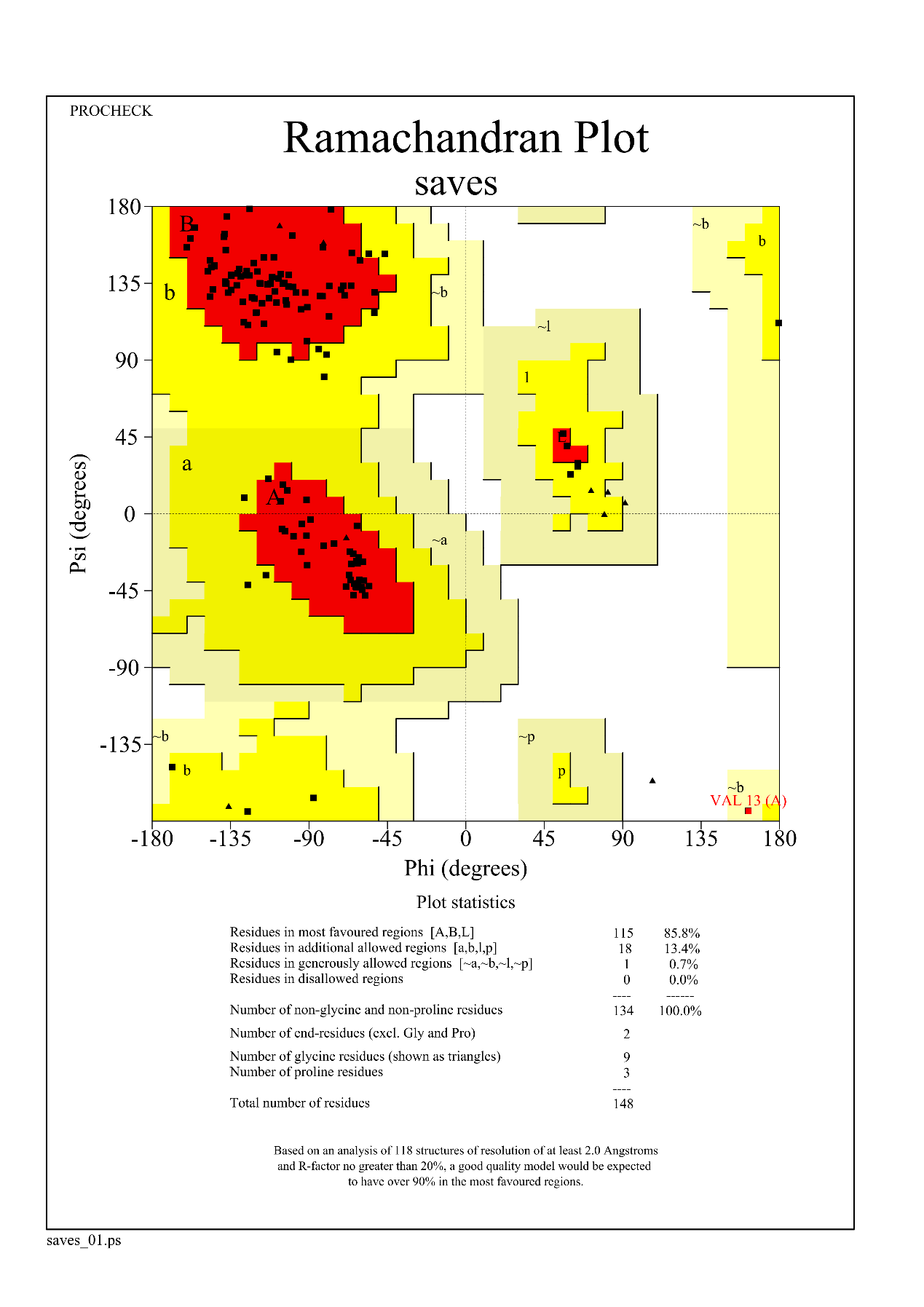

Fig. S4. Ramachandran plot of MjavOBP3 model.

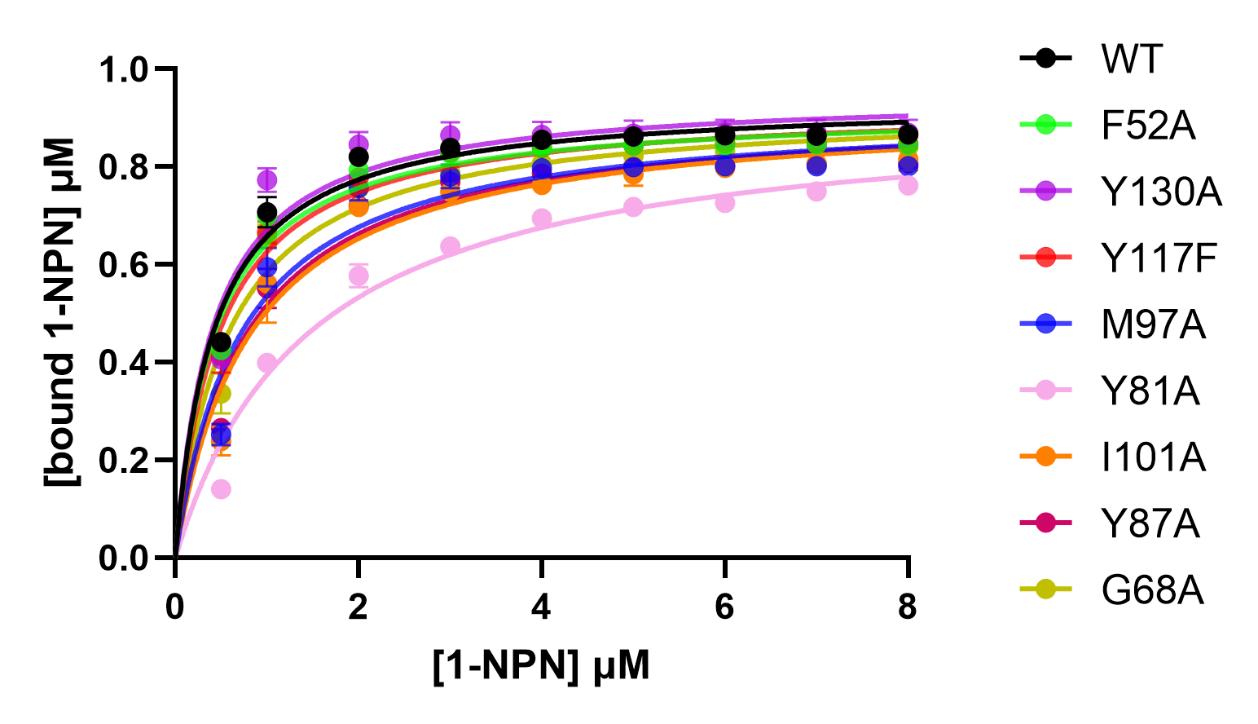

Fig. S5. Binding curve of 1-NPN to MjavOBP3 WT and mutants. The dissociation constants (K _d_) were 0.42±0.09 μM for WT, 0.43±0.08 μM for F52A, 0.58±0.14 μM for G68A, 1.46±0.38 μM for Y81A, 0.79±0.27 μM for Y87A, 0.72±0.21 μM for M97A ,0.83±0.27 μM for I101A, 0.48±0.11 μM for Y117F, 0.42±0.16 μM for Y130A,

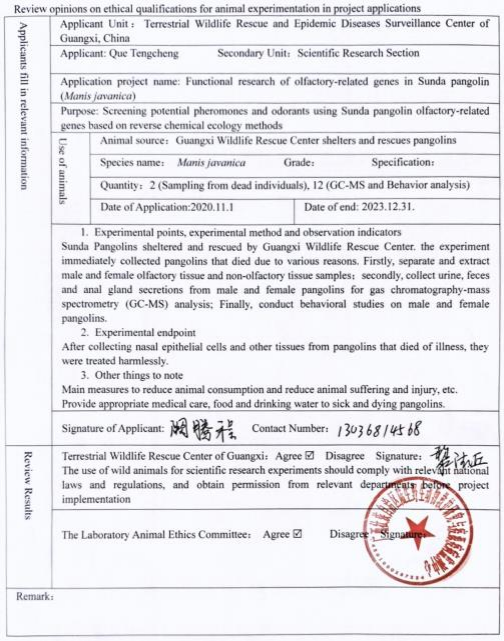

Fig. S6. Ethics statement
